# Prioritization of enhancer mutations by combining allele-specific chromatin accessibility with deep learning

**DOI:** 10.1101/2019.12.21.885806

**Authors:** Zeynep Kalender Atak, Ibrahim Ihsan Taskiran, Christopher Flerin, David Mauduit, Liesbeth Minnoye, Gert Hulsemans, Valerie Christiaens, Ghanem-Elias Ghanem, Jasper Wouters, Stein Aerts

**Affiliations:** VIB-KU Leuven Center for Brain & Disease Research, Leuven, Belgium; KU Leuven, Department of Human Genetics KU Leuven, Leuven, Belgium; Institut Jules Bordet, Université Libre de Bruxelles, Brussels, Belgium

## Abstract

Prioritization of non-coding genome variation benefits from explainable AI to predict and interpret the impact of a mutation on gene regulation. Here we apply a specialized deep learning model to phased melanoma genomes and identify functional enhancer mutations with allelic imbalance of chromatin accessibility and gene expression.

## Main

Genomic sequence variation within enhancers and promoters can have a significant impact on the cellular state and phenotype, as illustrated by many disease-associated SNPs^1^ and mutations^2^ in the non-coding genome. However, sifting through the millions of candidate variants in a personal genome or cancer genome, to identify only those variants that impact enhancer function, remains a major challenge. Commonly, two approaches are taken to overcome/tackle this problem. The first one is quantitative trait loci (QTL) analysis in which a variant is correlated to a cellular trait across a large number of samples. This strategy is widely used with expression data, as well as chromatin immunoprecipitation and chromatin accessibility data, for detecting QTL associated with gene expression (eQTL), transcription factor binding (bQTL)^3^, histone modifications (hQTL)^4^, or chromatin accessibility (caQTL)^5^ respectively. This type of analysis suffers from the requirement of large sample sizes, since the effect sizes of these variants are usually low. The alternative approach is to assess allelic imbalance at a heterozygous site itself by comparing the two alleles of the diploid genome against each other. This allele counting approach is extensively used with RNA-seq data to identify allele-specific expression variants^8^, but is also applicable to other types of functional genomics data^6^. Here, the strategy relies on finding the allelic origin of the signal observed in functional genomics data. The use of a personalized diploid genome prevents potential technical biases and ensures obtaining accurate allelic counts^6,7^. Importantly, this approach can work even with a single sample, if (phased) high quality variants are available and a personalized diploid genome can be reliably constructed. This is especially true for cancer genomes that often have a large number of differences from the reference genome.

In this study we investigate how allele-specific changes in chromatin accessibility - as a proxy for transcription factor occupancy - combined with deep learning and motif analysis can be exploited to distinguish functional from non-functional enhancer mutations in phased melanoma genomes (**Figure 1a**). We obtained haplotype-resolved whole genome sequence (WGS) data of 10 patient-derived melanoma cultures (MM lines) using linked-reads technology from 10X Genomics. Each sample was sequenced to an average depth of 38X (except MM087 at 68X and MM099 at 133X). We also obtained chromatin accessibility profiles of the same melanoma lines with ATAC-seq^8^ (9 samples were profiled with OmniATAC-seq and were obtained from Wouters *et al* ^8^; while ATAC-seq data for A375 was downloaded from GSE82330^9^). For combining the personal diploid genomes with ATAC-seq, we used a modified *alleleseq* pipeline to obtain allele-specific chromatin accessibility variants (ASCAVs)^10^ (**Figure 1a,b**) (see **Materials and Methods**). The BaalChIP^11^ method then enabled us to correct the observed allelic ratios from ATAC-seq reads using the genomic allelic ratios from the whole genome sequence data to correct for copy number variations that are frequently observed in these lines (**Figure S1**). From a total of 16 million phased variants across 10 genomes (**Figure 1c**), 139,420 variants overlap with ATAC-seq peaks, from which we identified 20,464 (14.7%) to represent significant ASCAVs across the 10 genomes (ranging from 451 to 7,183 per sample) (**Table S3,4**). The majority of the ASCAVs are unique to one MM line, and a small proportion shared between multiple samples (1,533 out of 20,464; 7.5%) (**Figure S2**). Only two of the shared ASCAVs are called discordantly between the samples (rs61794783 with G>C/T, and rs673002 with G>T/C) which are multiallelic sites (dbSNP150), illustrating the accuracy of the ASCAV pipeline. We also assembled a set of control SNPs, namely heterozygous SNPs within ATAC-seq peaks that show no allelic bias. The genomic distribution of both sets, ASCAVs and control SNPs, is highly similar (**Figure S3, Table S5**).

**Figure 1.**
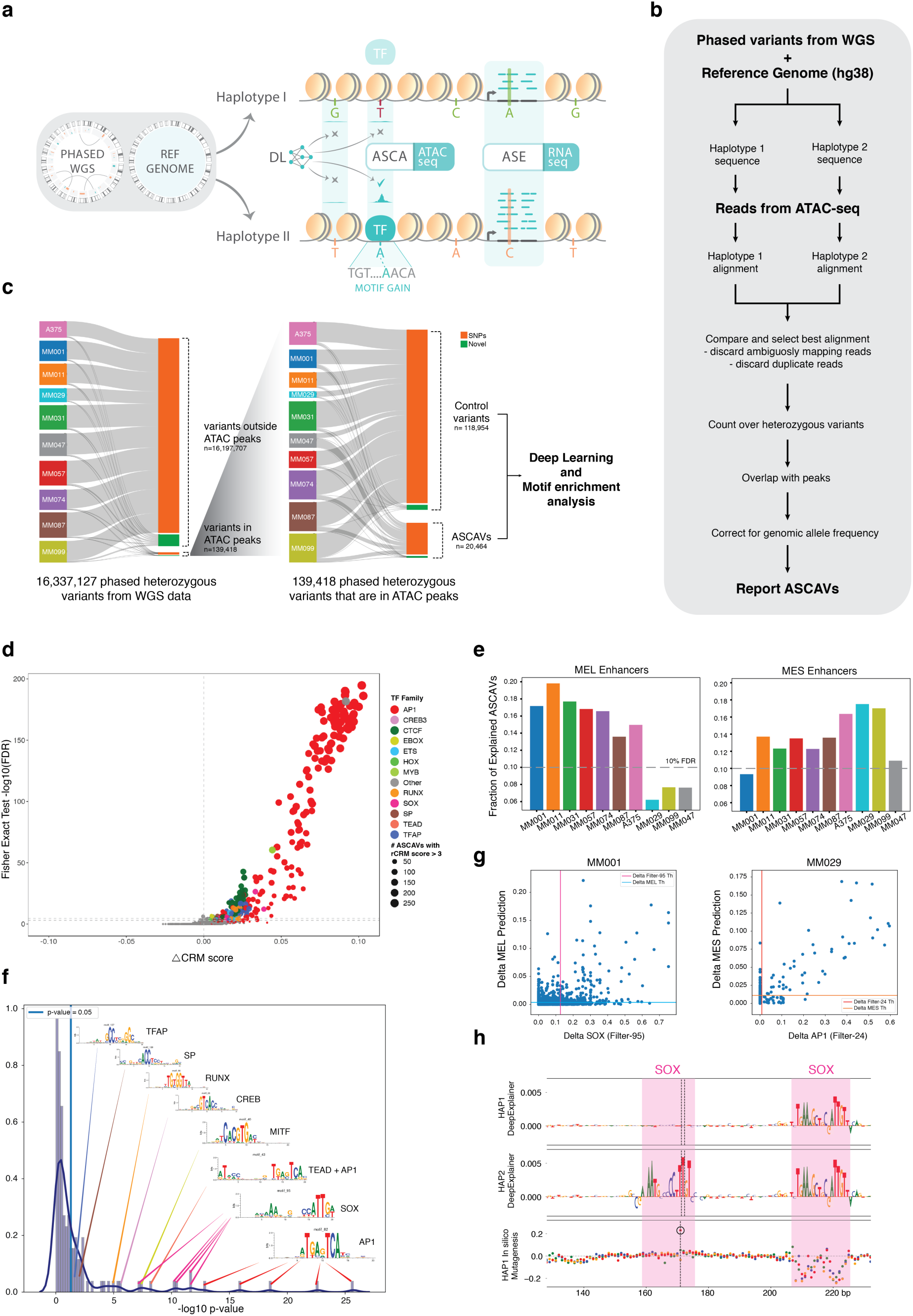
Detection of allele-specific chromatin accessibility. **a.** Phased whole genome sequencing is applied to 10 melanoma cell lines, and used together with the reference genome to create personalized diploid genomes. Matched ATAC-seq and RNA-seq data is used to detect allelic imbalance in chromatin accessibility (ASCA) or gene expression (ASE). By combining a melanoma-specific deep learning model (DeepMEL), and motif discovery, *cis*-regulatory variants are predicted. **b.** Analysis pipeline. **c.** Sankey diagram of the number of variants that went through our ASCA discovery pipeline. 16 million phased heterozygous variants across 10 melanoma genomes are filtered to retain 139,420 variants that overlap with an ATAC-seq peak, of which 20,464 present allelic imbalance. Novel/SNP ratio shows a significant shift in the ASCA variants compared to control variants (OR of 1.8 with FET p-value of <0.01) with an average 5.5% novel variants in ASCA group compared to 2.6 % novel variants in the non-ASCA group. **d.** Scatter plot of motifs that are associated (positively) with chromatin accessibility. The x-axis represents delta cluster-buster motif score (between preferred allele and reference), and the y-axis represents the -log scaled FDR corrected p-value. The sizes of the dots are proportional to the number of occurrences of the motif. Each motif is colored based on its direct or inferred similarity a transcription factor family. **e.** Fraction of DeepMEL explainable ASCAVs on MEL (left) and MES (right) enhancers for each MM line at 10% FDR. **f.** Density plot shows the p-value of each convolutional filter after FET analysis. The top filter that explains more fraction of ASCAVs is AP-1 and it is followed by SOX. **g.** Correlation plot of delta MEL prediction score vs delta SOX filter score (left) and delta MES prediction score vs delta AP-1 (right) caused by ASCAVs showing that SOX motif has a direct effect on MEL enhancer prediction while AP-1 motif has a direct effect on MES enhancer prediction. **h.** DeepExplainer plot of HAP1 (top) and HAP2 (middle). C>T ASCAV at position 171 generates a SOX binding site and it increases MEL enhancer score. *In silico* mutagenesis shows that only C>T increases the MEL enhancer score significantly. Mutations on the right SOX binding site decrease the MEL enhancer score.

When testing for allele-specific expression (ASE) using matched RNA-seq data on the same MM lines^12^ we identified 20,746 distinct ASE variants, associated with 9,787 genes (**Table S6**). One gene, namely *MAP2K3* (also known as MEK3), shows ASE in all 10 samples; this gene has been previously reported to be expressed in an allele-specific manner in various human and mouse tissues^13,14^. Globally, there is a significant enrichment of ASCAVs near genes with ASE (p-value 0.005 with Fisher’s exact test) with 12.8% of ASCAVs being located 2 kb upstream (promoter) or within an intron of an ASE gene (**Table S7**). However, despite this enrichment, using the proximity of ASE alone is not accurate enough to prioritize functional *cis*-regulatory variants.

A commonly applied strategy to predict the functional relevance of a putative *cis*-regulatory variant, is to predict its impact on the encompassing enhancer or promoter, namely the creation or destruction of a transcription factor binding site^2,5,15^. However, binding site predictions using position weight matrices (PWM) are notorious for their high false positive rates^16^. To overcome this problem, we propose a motif enrichment strategy to identify PWMs that present significantly more changes than expected by chance, across the entire set of ASCAVs. In addition, more advanced machine-learning models have been used that are trained on enhancers and take flanking sequence information into account, including gapped k-mer SVMs^17^ and deep learning models like Basset^18^, Basenji^19^, DanQ^20^, DeepBind^21^, DeepSea^22^, and FactorNet^23^. Such models are available in Kipoi^24^, and can be readily applied to score *cis*-regulatory variants. However, these generic models are usually trained on large collections of epigenomics data, allowing their employment to “any” cell type. In contrast, we follow a different strategy, specifically tailored towards the regulatory program of two melanoma cell states, namely the melanocytic state (MEL), and the mesenchymal-like state (MES)^8,12,25^. Particularly, in our companion paper (Minnoye & Taskiran et al., *companion paper submitted*), we trained a deep learning model, called DeepMEL, on a cohort of melanoma ATAC-seq data. We validated this model against melanoma ATAC-seq data from other species, as well as through functional enhancer-reporter assays. We have shown that DeepMEL outperforms other models in a benchmark study using saturation mutagenesis of the *IRF4* enhancer, measured by a massively parallel enhancer reporter assay^26^. DeepMEL is able to accurately predict the impact of enhancer mutations in the *IRF4* enhancer, and between species.

As a baseline experiment, before applying DeepMEL, we performed a motif enrichment analysis across the entire set of ASCAVs. By calculating a delta PWM score for each ASCAV (see **Materials and Methods**), for a collection of 22,000 PWMs^27^, we observed AP-1 motifs to be most significantly altered by ASCAVs compared to control SNPs, affecting 844 ATAC-seq peaks across all samples (overall 183 AP-1 family PWMs were enriched with FET adjusted p-value threshold of 0.05; **Figure 1d, Figure S4**). It has been previously reported that AP-1 can act as a pioneer factor that results in nucleosome displacement at enhancers in murine mammary epithelial cells^28^. A similar motif enrichment technique has been applied before to identify pioneer factors from chromatin accessibility QTL data^29^. These data suggest that occupancy of AP-1 is strongly correlated with changes in chromatin accessibility, and confirms the power of allele-specific chromatin accessibility to identify gain-of-function enhancer mutations. We also tested whether motif gains or losses can be negatively associated with accessibility, and interestingly identified only a few motifs from ZEB/SNAI family, which are known repressor transcription factors in the neural crest lineage cell types including in melanomas^30–33^ (**Figure S5**).

Next, we used DeepMEL to score all ASCAVs (**Materials and Methods**). We focused on the two best characterized enhancer classes in DeepMEL, namely MEL enhancers, which are governed by SOX10, TFAP2, MITF, and RUNX motifs; and MES enhancers governed by AP-1 and TEAD motifs. DeepMEL identifies significantly more changes to MEL enhancer sequences that correlate with chromatin accessibility, compared to the control SNPs that are not correlated with chromatin accessibility (**Methods, Figure 1e, Figure S6,S7**). Interestingly, explainable mutations in MEL enhancers occur more frequently in the samples of the melanocytic subtype, where they are operational. The MES enhancers, on the other hand, are mostly affected in samples of the mesenchymal subtype (**Figure 1e**). Note that, in agreement with the motif enrichment analysis, AP-1 motif changes (the main drivers of the MES scores, **Figure 2g**), are also found enriched for ASCAVs in the melanocytic lines, except for MM001 which has no AP-1 activity (**Figure S4**).

**Figure 2.**
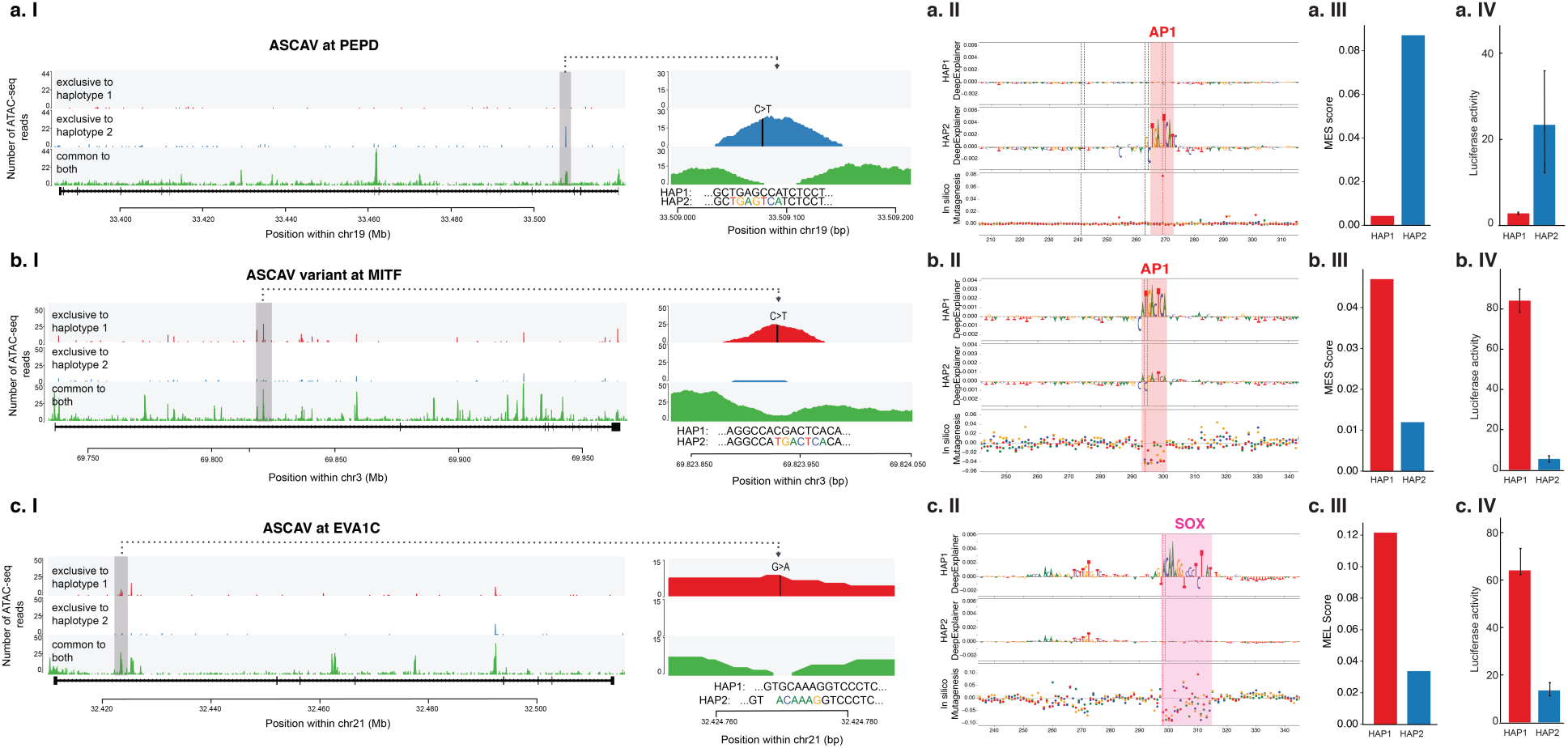
Explanation and validation of three cis-regulatory mutations. Each row (**a,b,c**) shows a detected ASCAV with in its locus with an inset of the allele-specific accessibility peak (**I**); the DeepExplainer plots and *in silico* mutagenesis of the two haplotypes (**II**); the DeepMEL score for both haplotypes (**III**); and the luciferase enhancer-reporter activity for both haplotypes (**IV**). **a.** C>T variant in *PEPD* intron generating an AP-1 site, with an increase in MES enhancer score. The *in silico* mutagenesis plot shows that only a single mutation to T at position 269 increases the MES enhancer prediction significantly, and this is exactly the location of the SNP. Control variants at position 241 and 263 do not change the enhancer score. **b.** ASCAV in an intron of *MITF* creating an AP-1 site. **c.** ASCAV in an intron of *EVA1C*, creating a SOX10 binding site.

The total number of ASCAVs that can be explained from each MM line by DeepMEL at 10% FDR for MEL and MES classes are 2,093 and 1,715 respectively (excluding MM047 ASCAVs). (**Methods, Figure S6,S7**). This percentage is higher compared to earlier studies^3,5,15,34–37^, where enhancer variants have been attributed to *cis*-regulatory changes in transcription factor binding sites. Next, we analyzed the function of these interpretable ASCAVs in more detail using several explainable AI (XAI) techniques. Firstly, we assessed which of the convolutional filters are mostly affected by ASCAVs, compared to control SNPs (**Figure 1f, Figure S8)**. In line with our previous observations, the AP-1 filter is the most frequently altered, followed by the SOX10. The second XAI technique we used is based on DeepExplainer^38^, which assesses how much each nucleotide in the enhancer contributes to its classification (**Figure 1h**). This analysis highlights the fact that enhancer mutations that affect chromatin accessibility mainly occur in TF binding sites within the enhancer, and that the most commonly affected motif in MEL enhancers is the SOX10 motif. Indeed, presence or absence of a SOX10 motif has the greatest influence on changes in ATAC-seq signal, among the MEL enhancer motifs (**Figure 1f, Figure S9**). Nevertheless, a subset of ASCAVs can also be explained by changes in TFAP2 and MITF motifs (**Figure S10, Figure S11**). Interestingly, and in line with our previous observations, SNPs that create a ZEB/SNAI motif result in a decrease in chromatin accessibility, suggesting that these factors promote nucleosome stability at the enhancer, or alternatively inhibit the nucleosome displacement effect of SOX10 or the nucleosome stabiliser effect of TFAP2 (Minnoye & Taskiran, *companion paper submitted*). Various members of the ZEB/SNAI family have indeed been suggested to act as transcriptional repressors^30–33^. Finally, a third technique we applied is *in silico* saturation mutagenesis whereby each possible mutation in an enhancer sequence is evaluated by re-classification (**Figure 1h)**. From this assay we find that ASCAVs overlap with *in silico* predicted vulnerable nucleotides (**Figure S10 and Figure S11**).

Our phased genomes allow linking the allelic imbalance of chromatin accessibility with the allele-specific expression of nearby genes. 12.6% of ASCAVs that can be explained by motif changes are located near (2 kb upstream or intronic) a gene that presents ASE. Therefore, these enhancer mutations may underlie the change in expression of the target gene. To further examine this, we selected three enhancers in MM057 for which the predicted target gene shows ASE. The first two examples, PEPD and MITF, have a predicted gain of an AP-1 motif, as indicated by DeepMEL (**Figure 2a,b, Figure S12)**. Luciferase reporter assays conducted in the same cell line using sequences of both haplotypes confirm the potency of these variants to drive enhancer activity, only when the AP-1 site is present (**Figure 2a,b**). The third example is an enhancer in the first intron of EVA1C with a predicted SOX10 motif gain by the DeepMEL model. This variant (rs2833812) is identified as a phased heterozygous SNP in four lines (MM031, MM057. MM074, MM087) and results in allele specific accessibility in all cases (**Figure S13**). Again, when assessed with luciferase reporter assay (in MM057), only the enhancer sequence that carries the SNP generating the SOX10 motif is able to drive luciferase activity (**Figure 2c**). This suggests that enhancer mutations associated with changes in chromatin accessibility can indeed have an effect on the expression of nearby target genes.

In conclusion, with a specialized deep learning model that is trained and validated on a cohort of melanoma cultures (Minnoye & Taskiran, *companion paper submitted*), we detected nearly five thousand variants across 10 samples. These variants show allele-specific chromatin accessibility and can be explained by mutations causing gains or losses of transcription factor binding sites, where certain factors have a stronger effect on changes in accessibility, including AP-1 in mesenchymal enhancers, and SOX10 in melanocytic enhancers. Although it is likely that a large fraction of these events mark *de novo* binding of a TF, without any consequence on gene expression, a fraction of these also impact enhancer activity and are associated with changes in gene expression. Our approach is applicable to any cancer type and may contribute to the identification of *cis*-regulatory driver mutations.

## Materials and Methods

### Cell culture

The melanoma MM lines are derived from patient biopsies by the Laboratory of Oncology and Experimental Surgery (Prof. Dr. Ghanem Ghanem) at the Institut Jules Bordet, Brussels^8,12,39^. Cells were cultured in Ham’s F10 nutrient mix (ThermoFisher Scientific) supplemented with 10% fetal bovine serum (Invitrogen) and 100 µg/ml penicillin/streptomycin (ThermoFisher Scientific). The commonly-used, human melanoma cell line A375 was obtained from the ATCC and was maintained in Dulbecco’s Modified Eagle’s Medium with high glucose and glutamax (ThermoFisher Scientific), supplemented with 10% fetal bovine serum and penicillin-streptomycin. Cell cultures were kept at 37°C, with 5% CO2 and were regularly tested for mycoplasma contamination, and were found negative. For each cell line, 1 million cells were used as input for the extraction of genomic DNA.

### Phased whole genome library preparation

The extraction of high molecular weight (HMW) genomic DNA (gDNA) and subsequent preparation of phased whole genome libraries was performed using the Chromium instrument and the Linked-Reads Genome Kit v2 (10x Genomics), according to the manufacturer’s protocol (Rev A). Briefly, first, HMW gDNA was isolated using the MagAttract HMW DNA Kit (Qiagen). Quality of the HMW gDNA (length and integrity) was verified using the Genomic DNA ScreenTape on the Tapestation instrument (Agilent). Next, the HMW gDNA was carefully, serially diluted to final concentrations of 0.8 – 1.0 ng/μl (measured using the Qubit High Sensitivity kit), before loading onto the microfluidic Genome Chip (10x Genomics). After generation of nanoliter-scale Gel bead-in-EMulsions (GEMs), genomic fragments were barcoded during isothermal incubation at 30C in a C1000 Touch Thermal Cycler (Bio Rad). After incubation, the droplets were broken and the pooled barcoded fragments cleaned with a Cleanup Mix containing DynaBeads (Thermo Fisher Scientific). Next, the fragments were end-repaired, A-tailed and index adaptor ligated, with cleanup in between steps using SPRIselect Reagent Kit (Beckman Coulter). The Post-ligation product was then amplified with a C1000 Touch Thermal Cycler during the Sample Index PCR. Finally, the sequencing-ready library was cleaned up with SPRIselect beads. Libraries were sequenced on a NovaSeq6000 S2 flow cell with paired-end 2 × 150 cycli.

Sequence reads were processed using 10X Genomics companion software longranger (v2.2.2). Short variant calling was performed with GATK (v3.5) using --vcmode gatk option in longranger. Structural variants were called using Sniffles (v1.0.10). Single nucleotide variants and indels were further filtered with depth of sequencing filter of 20.

### Personalized genome construction

Variant calls (as generated by longranger) and structural variation calls (as generated by Sniffles) were used to construct personalized genomes with Crossstich. This procedure resulted in generation of personalized reference sequences per haplotype in fasta format as well as chain files that link reference genome to personalized genomes. To obtain chain files to link personalized genomes to reference genome, we performed whole genome alignment between personalized genomes and reference genome using Blat. Briefly, we partitioned the reference genome into 3kb pieces per chromosome and mapped to personalized genomes with Blat using the following options: -tileSize=11 - fastMap -minIdentity=95 -noHead -minScore=100. Resulting alignment files in psl format was combined, and chained with axtChain command and then sorted with chainMergeSort command. Then, alignment nets were created from chain files using chainNet command, and finally liftOver chain files were created with netChainSubset command.

### ATAC-seq library preparation

ATAC-seq data was generated using the OmniATAC-seq technique as described previously^40^. Cells were washed, trypsinized, spun down at 1000 RPM for 5 min, medium was removed and the cells were resuspended in 1 mL medium. Cells were counted and experiments were only continued when a viability of above 90% was observed. 50,000 cells were pelleted at 500 RCF at 4°C for 5 min, medium was carefully aspirated and the cells were washed and lysed using 50 uL of cold ATAC-Resupension Buffer (RSB) (see Corces et al., 2017 for composition) containing 0.1% NP40, 0.1% Tween-20 and 0.01% digitonin by pipetting up and down three times and incubating the cells on ice for 3 min. 1 mL of cold ATAC-RSB containing 0.1% Tween-20 was added and the eppendorf was inverted three times. Nuclei were pelleted at 500 RCF for 10 min at 4°C, the supernatant was carefully removed and nuclei were resuspended in 50 uL of transposition mixture (25 uL 2x TD buffer (see Corces et al., 2017 for composition), 2.5 uL transposase (100 nM), 16.5 uL DPBS, 0.5 uL 1% digitonin, 0.5 uL 10% Tween-20, 5 uL H2O) by pipetting six times up and down, followed by 30 minutes incubation at 37°C at 1000 RPM mixing rate. After MinElute clean-up and elution in 21 uL elution buffer, the transposed fragments were pre-amplified with Nextera primers by mixing 20 uL of transposed sample, 2.5 uL of both forward and reverse primers (25 uM) and 25 uL of 2x NEBNext Master Mix (program: 72°C for 5 min, 98°C for 30 sec and 5 cycles of [98°C for 10 sec, 63 °C for 30 sec, 72°C for 1 min] and hold at 4°C). To determine the required number of additional PCR cycles, a qPCR was performed (see Buenrostro et al., 2015 for the determination of the number of extra cycles). The final amplification was done with the additional number of cycles, samples were cleaned-up by MinElute and libraries were prepared using the KAPA Library Quantification Kit as previously described^40^. Samples were sequenced on a HiSeq4000 or NextSeq500 High Output chip.

### ATAC-seq alignment to the reference and personalized genomes

We aimed at obtaining minimum 15M usable reads per sample, and eventually achieved 41M reads on average across 9 sequenced samples (see Methods) (**Table S1**). Paired-end reads were mapped to the human genome (hg38) using bowtie2 with --very-sensitive option (v2.2.6). Mapped reads were sorted using SAMtools (v1.8) and duplicates were removed using Picard MarkDuplicates (v1.134). Reads were filtered by removing chromosome M reads and filtering for Q>2 using SAMtools. Usable reads is defined as the number of reads retained after these filtering steps. Paired-end reads were mapped to the sample specific personalized genomes using bowtie2 (v2.2.6). Mapped reads were sorted using SAMtools (v1.8) and duplicates were removed using Picard MarkDuplicates (v1.134). Reads were filtered by removing chromosome M reads and filtering for Q>2 using SAMtools. We observed that the number of reads mapping to personalized genomes were slightly higher than the number of reads mapping to the reference genome (hg38), which has been reported previously for ChIP-seq data^6,41^ (**Table S2**).

### ATAC-seq peak calling

Peaks from ATAC-seq data was called for reference mapped and personalize genome mapped data using MACS2 (v2.1.2) using the parameters -q 0.05, --nomodel, --call-summits, --shift -75 --keep-dup all and --extsize 150. Summits for personalized genome mapped samples were lifted over to reference genome using liftOver. Summits were extended by 250bp up- and downstream using slopBed (bedtools; v2.28.0), providing human chromosome sizes and filtered for blacklisted regions of the reference genome (ENCSR636HFF). Per sample, we obtained reference mapped peaks, HAP1 mapped peaks and HAP2 mapped peaks. To obtain a consolidated peak set per sample, we followed the strategy described by Corces et al^42^. Briefly, peak scores within each peakset was standardized by dividing peak scores to the sum of all peak scores for that sample. Haplotype 1 and 2 mapped peak sets lifted over to reference genome and consolidated peak set per sample obtained by an iterative filtering strategy: peaks were ranked by their normalized peak score, and any peak that overlapped with the highest scoring peak was filtered out. Next, the second highest scoring peak is processed this way, and the same procedure is repeated until a non-overlapping peak set is obtained. This strategy enabled us to obtain 500bp fixed-width peaks that are not biased towards the reference genome (referred to as consolidated peak sets).

### Identification of allele-specific events in ATAC-seq data

We have built a new allele specific variant detection pipeline using the backbone of alleleSeq pipeline ^10^. We used Bowtie2 ^43^ to map ATAC-seq reads to personalized genomes as described above. Next, we marked duplicate reads in each alignment using Picard. Then, we evaluated two alignment files (ie. haplotype 1 mapped and haplotype 2 mapped) to identify the most likely origin of each read. Each read was evaluated iteratively using mapping quality (MapQ), CIGAR string and XM tag (which reports the number of mismatches in the alignment) If the read had the same mapQ, same CIGAR string (or the same number of Ms) and same XM tag for both alignments, it was marked as commonly mapping. This step resulted in four bam files: haplotype1.exclusive, haplotype2.exclusive, haplotype1.common and haplotype2.common.

After identifying the source of each read, we filtered out duplicate reads. We also filtered out ambiguously mapping reads by evaluating the reads that map equally well to both haplotypes (ie. reads in haplotype1.common and haplotype2.common alignment files) in the reference genome. We lifted these positions over to hg38 coordinates, and discarded reads if it mapped to different locations. Next, we overlapped phased heterozygous variants obtained from whole genome sequence data with consolidated peak set (as described above). The variant positions that overlapped with the peaks were lifted over to haplotype 1 and 2 coordinates, and allele counts were obtained using samtools mpileup command with all four alignment files (allelic counts coming from common alignment files were compared and no major differences were found). Then allelic counts over heterozygous sites were merged, and variants that had at least 6 reads were further processed for allele specific accessibility analysis with BaalChIP^11^ package in R/Bioconductor. Count tables containing number of reference and alternative supporting reads per variant, together with allelic ratio of the same variant from whole genome sequence data was provided to runBayes command of BaalChIP, and allele-specific chromatin accessibility variants were identified. The remaining variants were defined as control variants.

Genomic annotations of both sets of variants were done using ChIPSeeker with UCSC hg38 knownGene table (TxDb.Hsapiens.UCSC.hg38.knownGene package in R/Bioconductor).

### Motif enrichment analysis with allele-specific binding events

ASCA and control variants per sample were overlapped with consolidated ATAC-seq peaks. Peaks that had multiple variants were filtered out if the allelic bias between variants was inconsistent. Allelic counts were used to determine preferred allele (i.e. allele that has the highest ATAC-seq signal). Peak sequences were extracted from the fasta sequence of the preferred allele (using fastaFromBed command from bedtools^44^) Reference sequence for each variant was extracted from hg38 using the same command. For each peak, we calculated the CRM score for the preferred allele and the other allele (reference sequence) using a set of 22,000 position weight matrices ^27^, and calculated a “delta CRM score” for each peak and for each motif (**Figure 1d**). We evaluated the enrichment of CRM delta’s in ATAC-seq peaks using one-sided Fishers’ exact test with a control set of 82,536 peaks containing non-ASCA variants. We performed enrichment analysis individually (per MM-line) and globally (across all MM-lines) (**Figure 1d, Figure S4**). Haplotype resolved ATAC-seq alignment figures were created with fluff ^45^.

### Identification of allele-specific expression variants

RNA-seq reads were mapped personalized genomes using Bowtie2 ^43^ with *--very-sensitive* option. We implemented the same post-processing steps as in ATAC-seq analysis; this included choosing the best alignment between two mappings based on mapping quality and number of mismatches, as well as removal of ambiguously mapping or duplicate reads. Next, we overlapped phased heterozygous variants obtained from whole genome sequence data with coding genome (hg38 CDS regions). Variant positions that overlapped with genes were lifted over to haplotype 1 and 2 coordinates, and allele counts were obtained using samtools mpileup command. Then allelic counts over heterozygous sites were merged, and variants that had at least 10 reads were further processed for allele specific expression variant analysis with binomial testing. Multiple testing correction was implemented with Benjamini & Hochberg method, and variants with FDR < 0.05 was reported as allele specific expression variants.

### The DeepMEL neural network

In our companion paper we describe DeepMEL (Minnoye & Taskiran et al., *submitted*). Briefly, DeepMEL is a hybrid CNN-RNN deep learning enhancer classification model trained on melanoma-specific co-accessible region classes. DeepMEL takes a 500bp DNA sequence and predicts an output vector corresponding to 24 classes. The model identifies TF binding sites on an enhancer and predicts the effect of a single mutation on the melanoma-specific enhancer identity, accessibility, and activity.

### Scoring enhancers and ASCAVs with DeepMEL

To score ASCAVs, we perturbed the 500bp ATAC-seq peaks by doing a single nucleotide change according to variants coming from two alleles. We calculated delta prediction score for each of the ASCAVs and the control variants for each of the classes. Then, we evaluated the delta prediction scores for each class (mainly MEL and MES classes) to identify the fraction of explainable ASCAVs using a one-sided Fishers’ exact test with a control set of 118,954 non-ASCA variants at 10% FDR.

### Assessing the mostly affected convolutional filters

To identify the most affected convolutional filters (128 filters in the convolutional layer of DeepMEL) by ASCAVs, we calculated a “delta activation score” of each convolutional filter for each ASCAV individually by getting the output of the convolutional layer using the enhancer sequence as input, before and after creating a single nucleotide change according to the ASCAV. We evaluated the enrichment of the delta activation scores for each filter using a one-sided Fishers’ exact test with a control set of 118,954 non-ASCA variants at 10% FDR.

### Calculating the contribution of each nucleotide to the final output

We initialised DeepExplainer ^38^ with randomly selected sequences (500) and calculated the importance scores of the sequence of interest with respect to any of the 24 classes. We multiplied this importance score by the one-hot encoded matrix of the sequence. Finally, we visualised the sequence by adjusting the nucleotide heights based on their importance score, similar to earlier work ^46^.

### In silico saturation mutagenesis

For a 500bp sequence, we generated mutated sequences by changing each single nucleotide into the three other possible nucleotides. We scored the initial sequence without mutations, as well as all 1500 generated sequences with DeepMEL and calculated the delta prediction score for each class and for each mutation by comparing the final prediction relative to prediction for the initial sequence.

### Luciferase assays

501 bp regions, surrounded by 20 bp flanking adaptors, were synthesis (TWIST Bioscience) and then individually cloned in a pGL4.23 plasmid. Luciferase activity in MM057 was measured using the Dual-luciferase Reporter Assay System (Promega). Briefly, 125,000 cells were seeded per well of a 24 well plate. The following day, cells were transfected with 40 ng renilla plasmid + 400 ng pGL4.23 plasmid using Lipofectamine 2000 (Invitrogen). 48 hours following transfection, cell lysates were prepared with Passive Lysis Buffer and Luciferase activity was measured on a VICTOR X3 plate reader (PerkinElmer). Luciferase assay values are represented in fold change of Fluc (pGL4.23 plasmid) vs Rluc (Renilla plasmid).

## Publicly available data used in this work

ATAC-seq data on A375 was obtained from GSE82330 ^9^. RNA-seq data (data for A375, MM001, MM011, MM029, MM031, MM047, MM057, MM074, MM087 and MM099) was downloaded from GSE60666 ^12^.

## Data and code availability

OmniATAC-seq data for the human lines MM001, MM011, MM029, MM031, MM047, MM074, MM057, MM087 and MM099 are available at the Gene Expression Omnibus, accession number GSE134432^8^. The whole genome sequence data generated for this study is being deposited to European Genome-phenome Archive (EGA; http://www.ebi.ac.uk/ega/). The code for the analysis of the WGS data and detection of ASCAV is deposited at https://github.com/aertslab/AS_variant_pipeline. The DeepMEL model is available from Kipoi.

## Acknowledgements

This work was supported by funded by an ERC Consolidator Grant to S.A. (no. 724226_cis-CONTROL), by the KU Leuven (grant no. C14/18/092 to S.A.), by the Foundation Against Cancer (grant no, 2016-070 to S.A.), a PhD fellowship from the FWO (L.M., no. 1S03317N) and a postdoctoral research fellowship from Kom op tegen Kanker (Stand up to Cancer), the Flemish Cancer Society, and from Stichting tegen Kanker (Foundation against Cancer), the Belgian Cancer Society (Z.K.A and J.W.). Computing was performed at the Vlaams Supercomputer Center and high-throughput sequencing via the Genomics Core Leuven. The funders had no role in study design, data collection and analysis, decision to publish or preparation of the manuscript.

## Author contributions

Z.K.A., I.I.T., J.W. and S.A. conceived the study. J.W., V.C., D.M., L.V.A. performed the experimental work for the whole genome sequencing and ATAC-seq. Z.K.A., C.F., G.H. performed the bioinformatics analyses. I.I.T. established the neural network and performed all analyses regarding the deep learning. Z.K.A., I.I.T., J.W., L.M. and S.A. wrote the manuscript.

