## Supplementary figures for "Prioritization of enhancer mutations by combining allele-specific chromatin accessibility with deep learning"

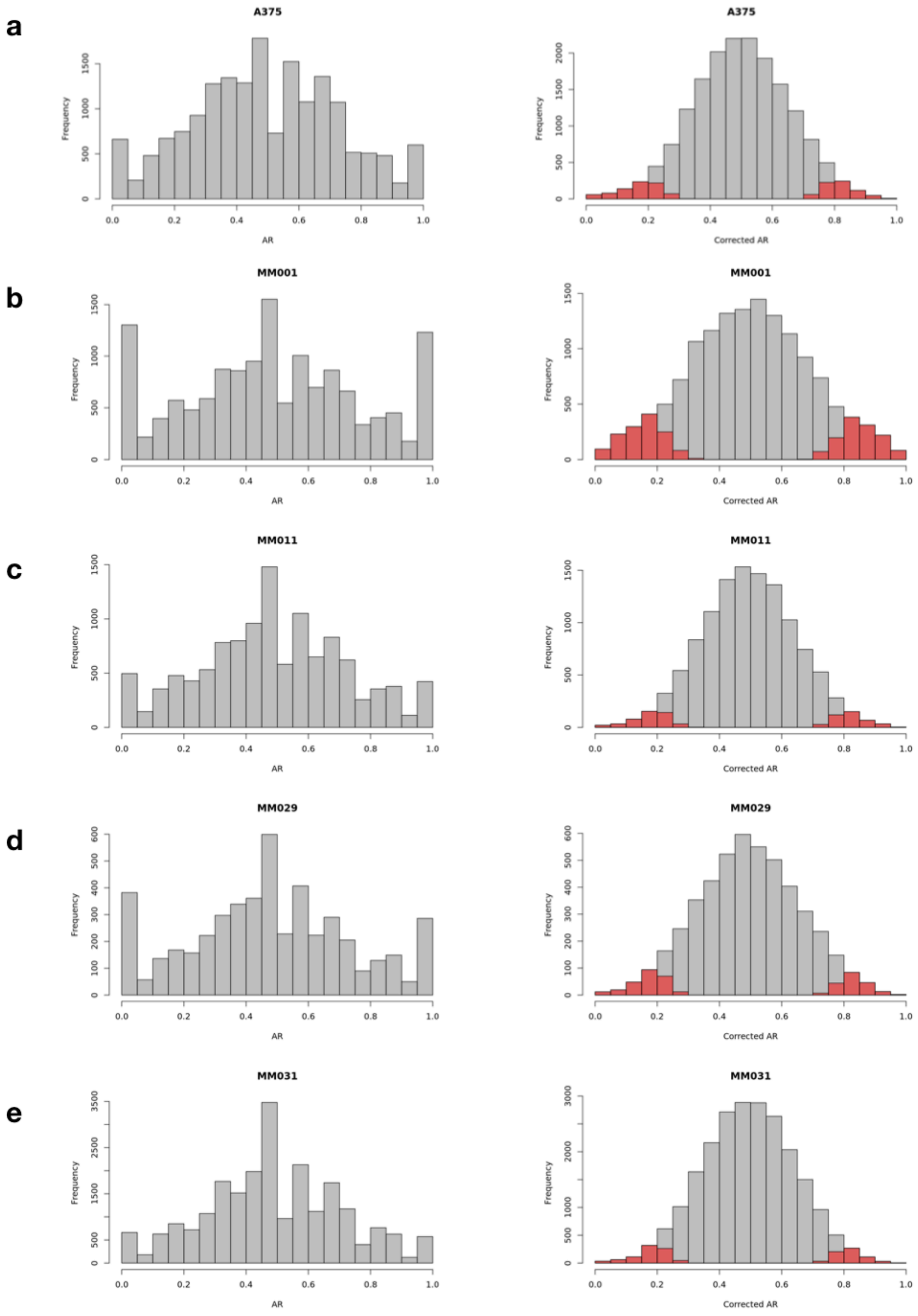

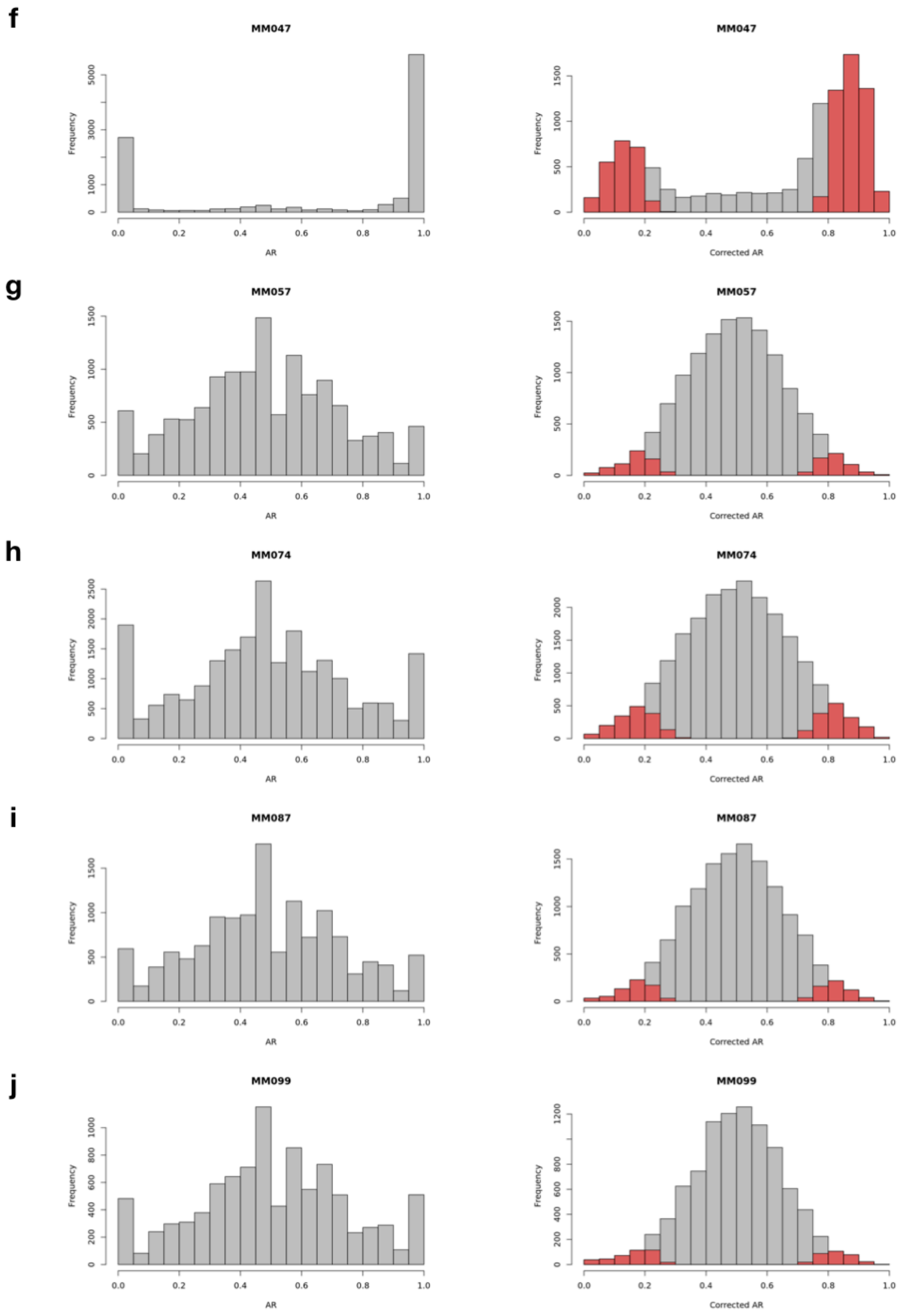


**Figure S1.** Histograms showing the distribution of allelic ratios (AR) before (on the left) and after (on the right) allelic ratio correction using BaalChIP. Corrected allelic ratios are estimated with BaalChIP using genomic allelic ratios from whole genome sequencing data. MM047 shows exceptionally high allelic ratios and is excluded from certain ASCAV analyses (see text).

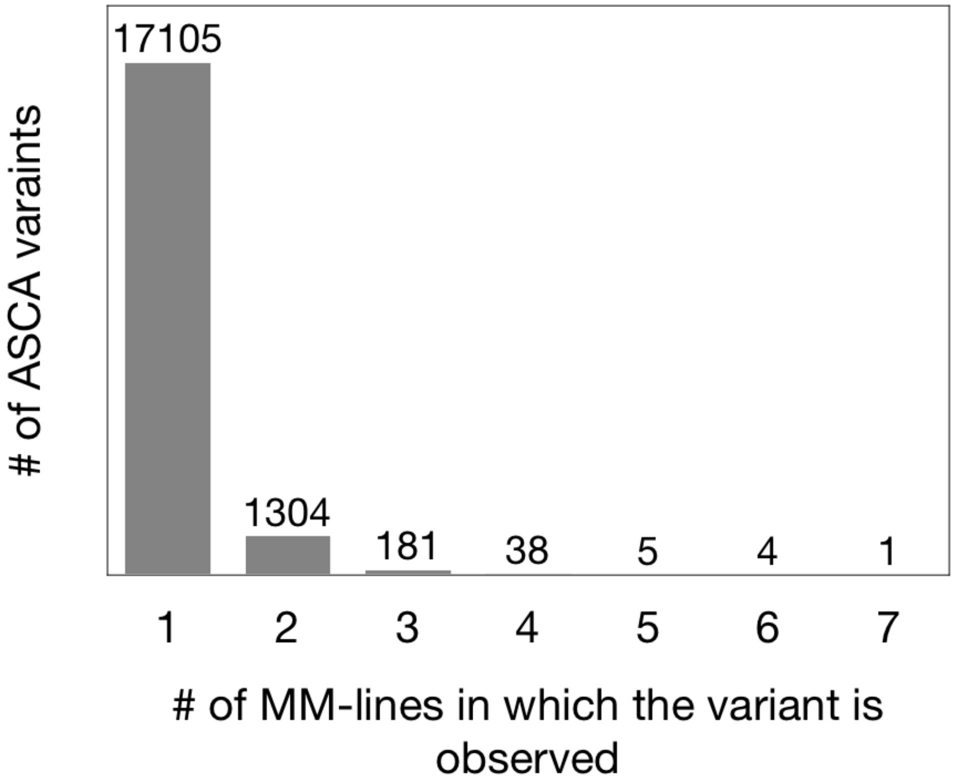


**Supplementary Figure 2.** Number of shared ASCA variants across ten MM-lines. Numbers above the bars indicate the number of variants shared between 2 to 7 MM-lines. 1,533 variants (1304+181+38+5+4+1) were detected in at least two MM-lines.

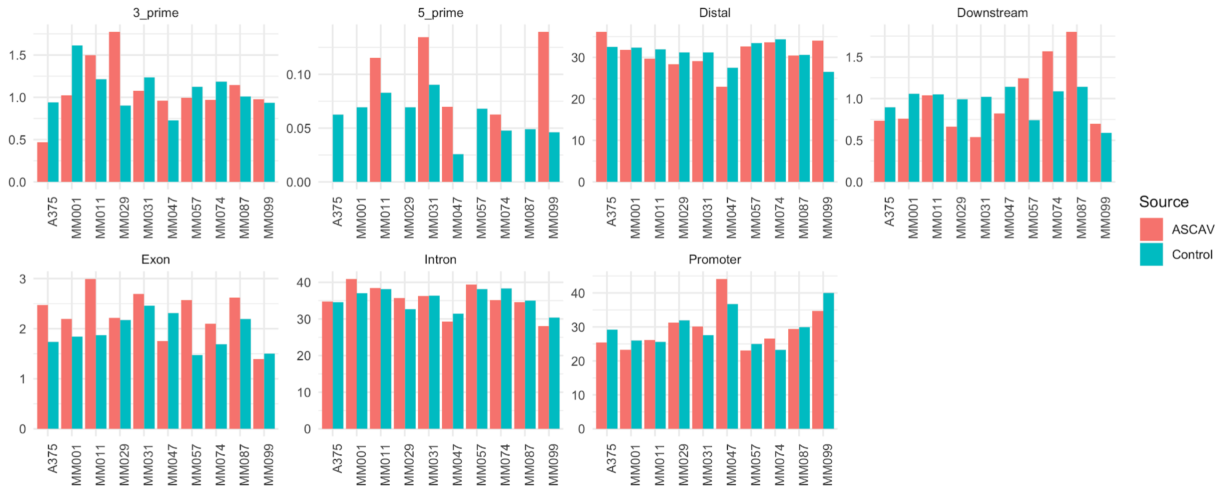


**Supplementary Figure 3.** Genomic localization of ASCA and control variants. Percentage of variants overlapping various genomic regions are presented in the barplot. ASCA and control variants do not display large differences in genomic localization.

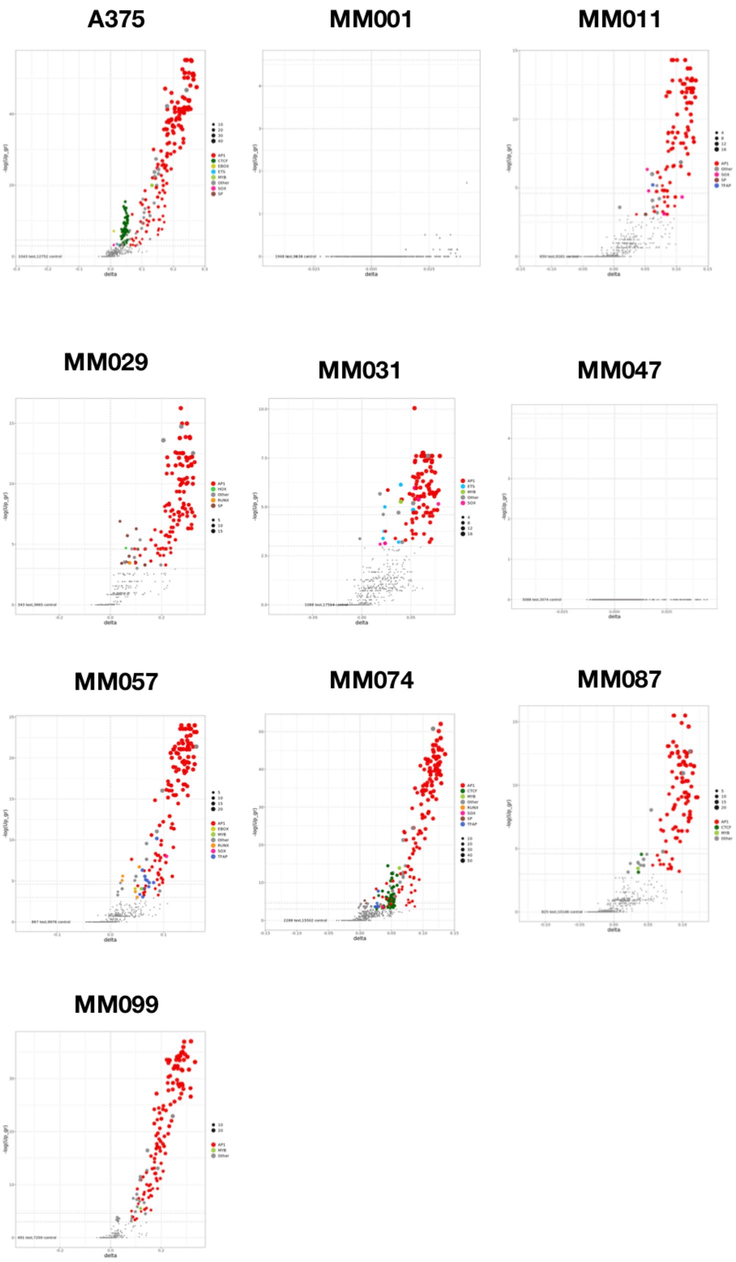


**Supplementary Figure 4.** Scatter plot of motifs that are associated (positively) with chromatin accessibility. Here the x-axis represents delta cluster-buster motif score, and the y-axis represents the -log10 scaled FDR corrected p-value (calculated by comparing ASCAVs with control SNPs using Fisher's Exact Test). The sizes of the dots are proportional to the number of occurrences of the motif. Each motif is colored based on its direct or inferred similarity a transcription factor family. MM001 and MM047 resulted in no significant motif enrichment using this analysis. MM001 is the only MM line without AP-1 activity, thus changes in AP-1 binding sites due to genomic variation have no effect on chromatin accessibility. MEL enhancers do show significant changes in MM001 (see text). MM047 presents too high allelic ratio (see Figure S1) to identify significant ASCAVs.

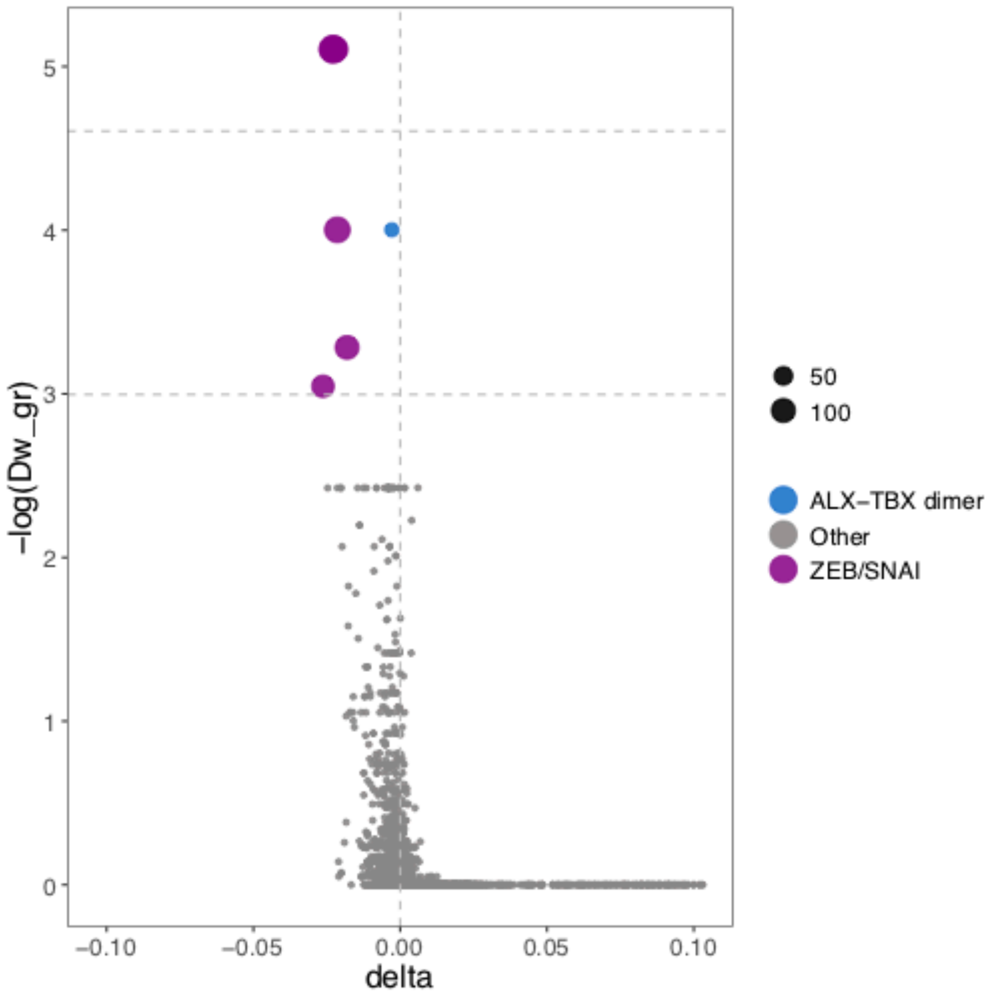


**Supplementary Figure 5.** Scatter plot of motifs that are associated (negatively) with chromatin accessibility. We tested whether motif gains or losses can be *discordant* with the allelic accessibility imbalance. In other words, a SNP that creates a TF binding site would cause a decrease in accessibility. Here, individual analysis per MM line resulted in no significant motif enrichment, however, due to the increased power in a global analysis across all samples, when all samples are merged, a few motifs showed enrichment, these are all from the ZEB/SNAI family, which are known repressor transcription factors in the neural crest lineage cells including melanomas[30]. These events are relatively rare and are only observed at 186 allelic imbalanced ATAC-seq peaks. DeepExplainer confirms this finding by identifying ZEB binding site gains associated with a decrease in accessibility on that allele (see Figure S10,S11 for examples).

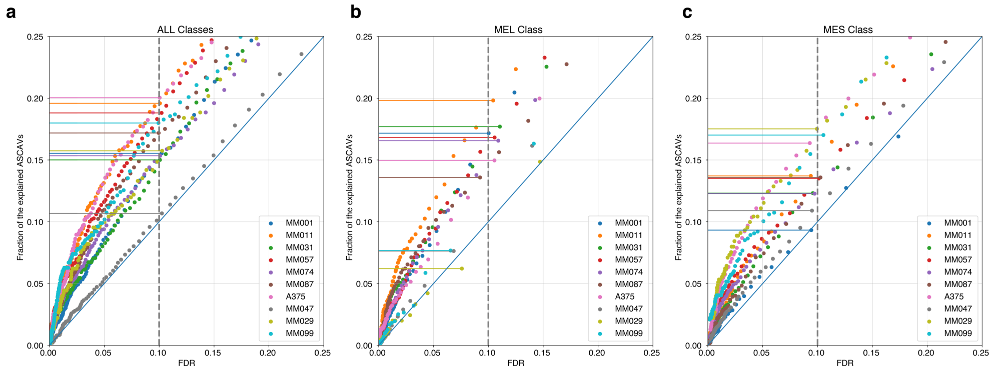


**Supplementary Figure 6. DeepMEL significance.**

**a.** Fraction of explainable ASCAVs when using maximum delta score among all classes. For all lines, the explained fraction ranges between 0.15 to 0.20 at 10% FDR (10% predicted by DeepMEL in the set control of SNPs). One sample, MM047 shows no enrichment, this is likely due to its high allelic ratios (see Figure S1). In total, 2,220 ASCAVs were explainable using the max delta across all classes (excluding MM047). **b.,c.** Instead of using the maximum delta score, ASCAVs are explained using each class separately, as illustrated for the two classes discussed in the main text, namely MEL (melanocytic enhancers), with 2,093 explained ASCAVs at 10% FDR; and MES (mesenchymal enhancers) with 1,715 MES ASCAVs explained (excluding MM047).

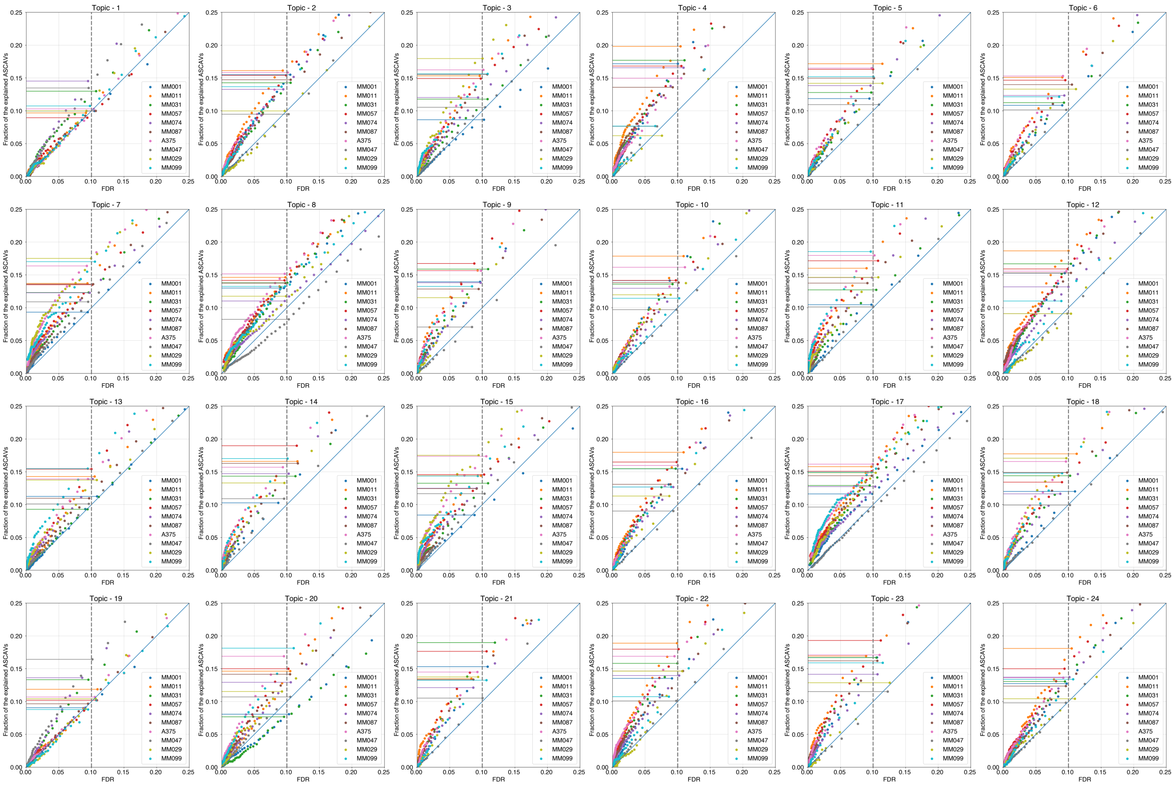


**Supplementary Figure 7.** DeepMEL was trained on 24 enhancer classes, each of them can be used to explain ASCAVs. We mainly focused on MEL (4th) and MES (7th) classes.

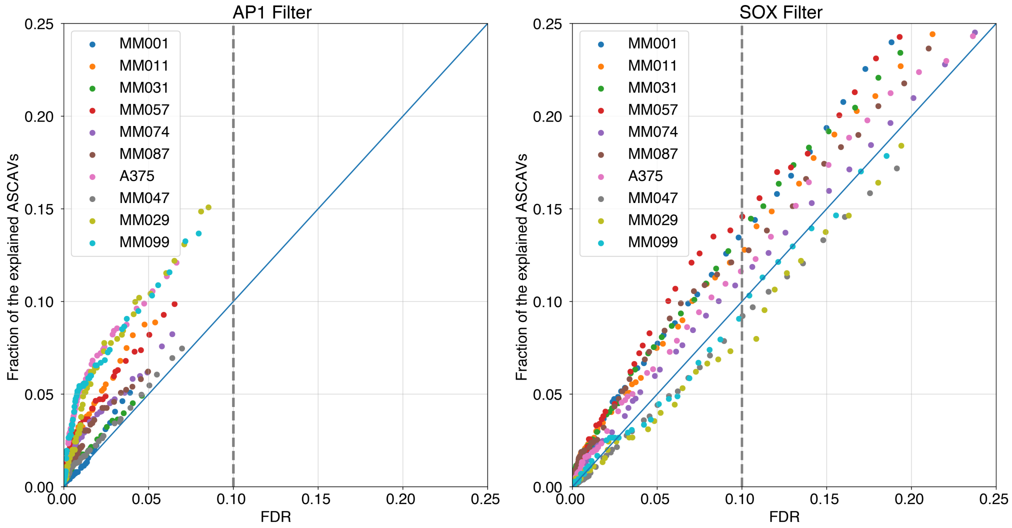


**Supplementary Figure 8.** Convolutional filters in the DeepMEL model can be used to test for motif enrichment. MM line specific motif enrichment is shown for the AP-1 (left) and SOX (right) filters. As expected, MES ASCAVs were explained better with the AP-1 filter in the mesenchymal-like cell lines (MM029 and MM099); while MEL ASCAVs are explained best with the SOX filter, in the melanocytic cell lines (MM001, MM031, MM057, MM074, MM087).

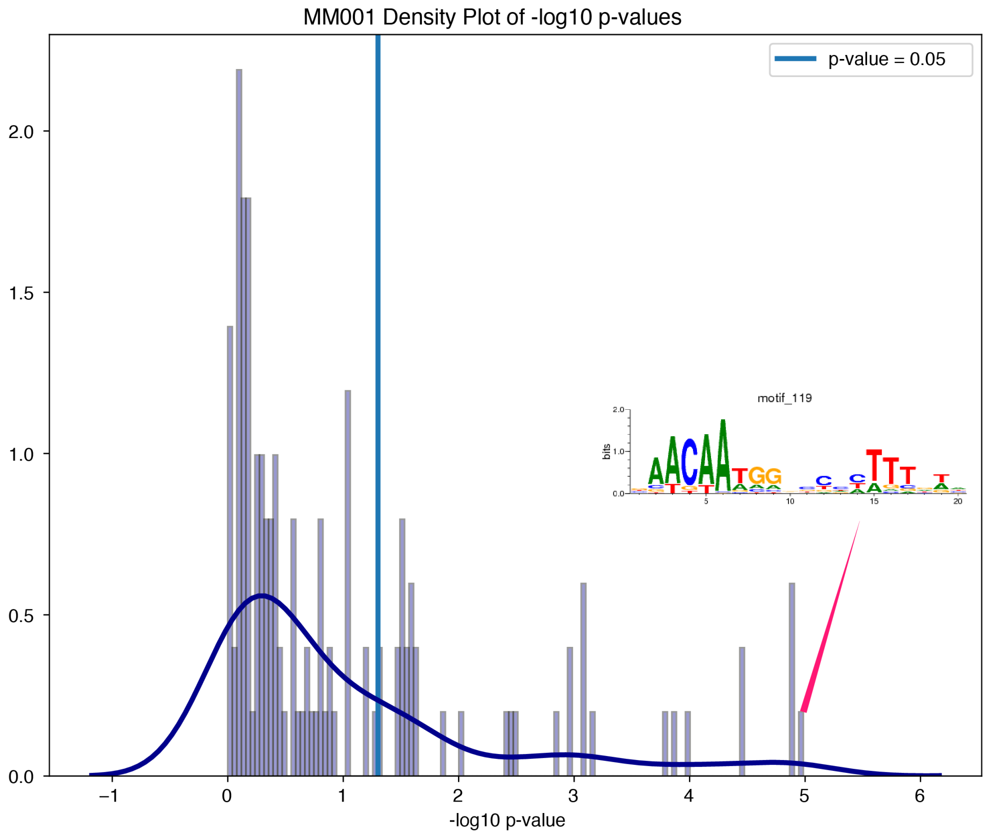


**Supplementary Figure 9.** Adjusted p-values (in -log10) resulting from a Fisher's Exact Test, comparing changes in scores of all convolutional filters in MM001 (a sample in the melanocytic state), for ASCAVs versus control SNPs, similar to Figure 1.f. Since MM001 has no AP-1 activity, the most important motif that explains ASCAVs is SOX.

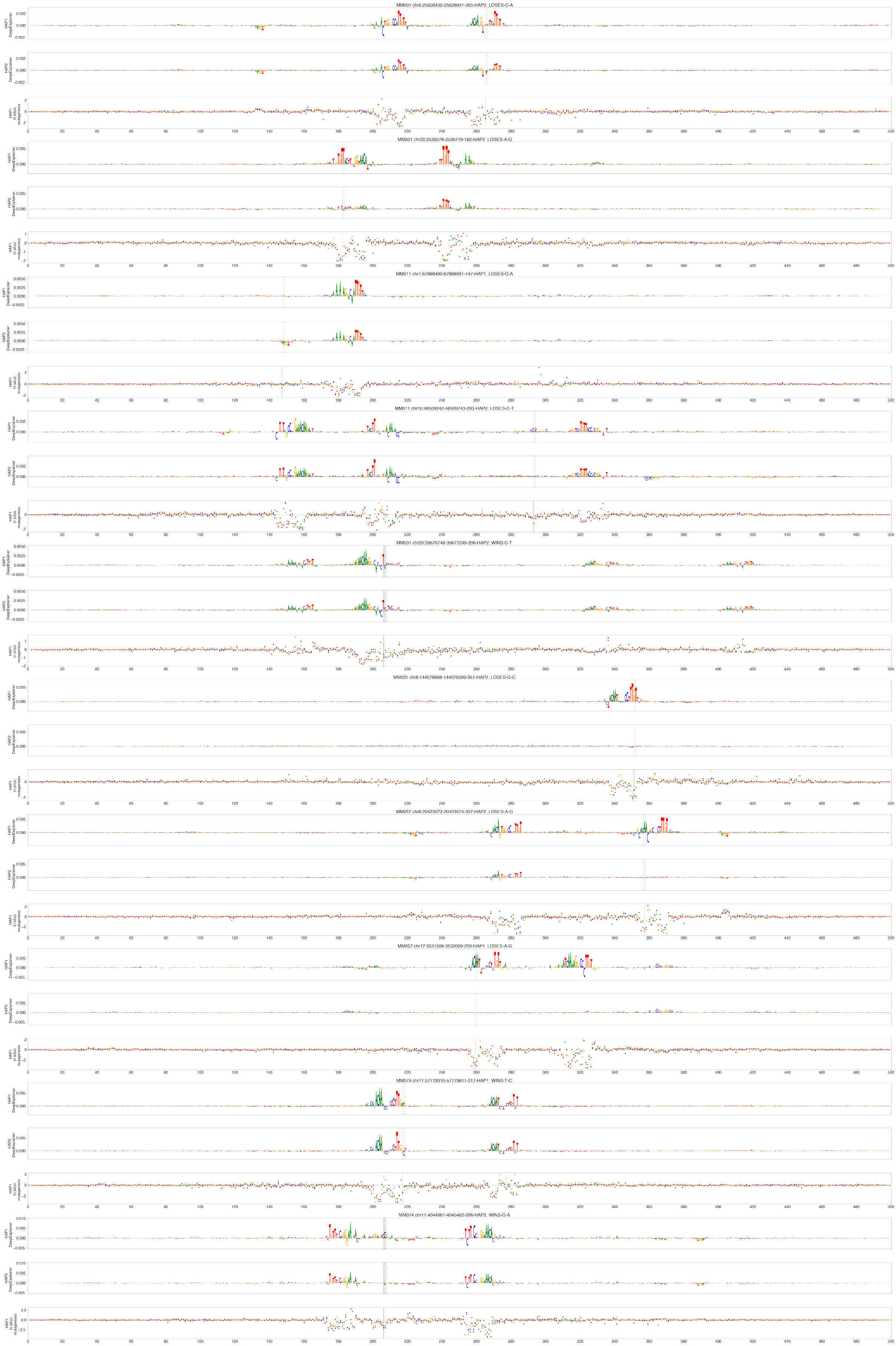
MM001 chr6:25628430-25628931-265-HAP2_LOSES-C-A
MM001 chr20:2539278-2539779-182-HAP2_LOSES-A-G
MM011 chr1:67888490-67888991-147-HAP1_LOSES-G-A
MM011 chr10:48509242-48509743-293-HAP2_LOSES-C-T
MM031 chr20:39676748-39677249-206-HAP2_WINS-C-T
MM031 chr8:144078898-144079399-351-HAP2_LOSES-G-C
MM057 chr8:20423073-20423574-357-HAP2_LOSES-A-G
MM057 chr17:3531508-3532009-259-HAP1_LOSES-A-G
MM074 chr11:57779310-57779811-217-HAP1_WINS-T-C
MM074 chr11:4044961-4045462-206-HAP2_WINS-G-A
HAP1 DeepExplainer
HAP2 DeepExplainer
HAP1 In silico mutagenesis

**Supplementary Figure 10.** DeepExplainer plots for several examples of MEL enhancers in 5 different MM lines (MM001, MM011, MM031, MM057, and MM074) with an ASCAV altering SOX (**i**, **iv**, **v**), TFAP (**ii**, **vi**), and ZEB (**ii**) binding sites.

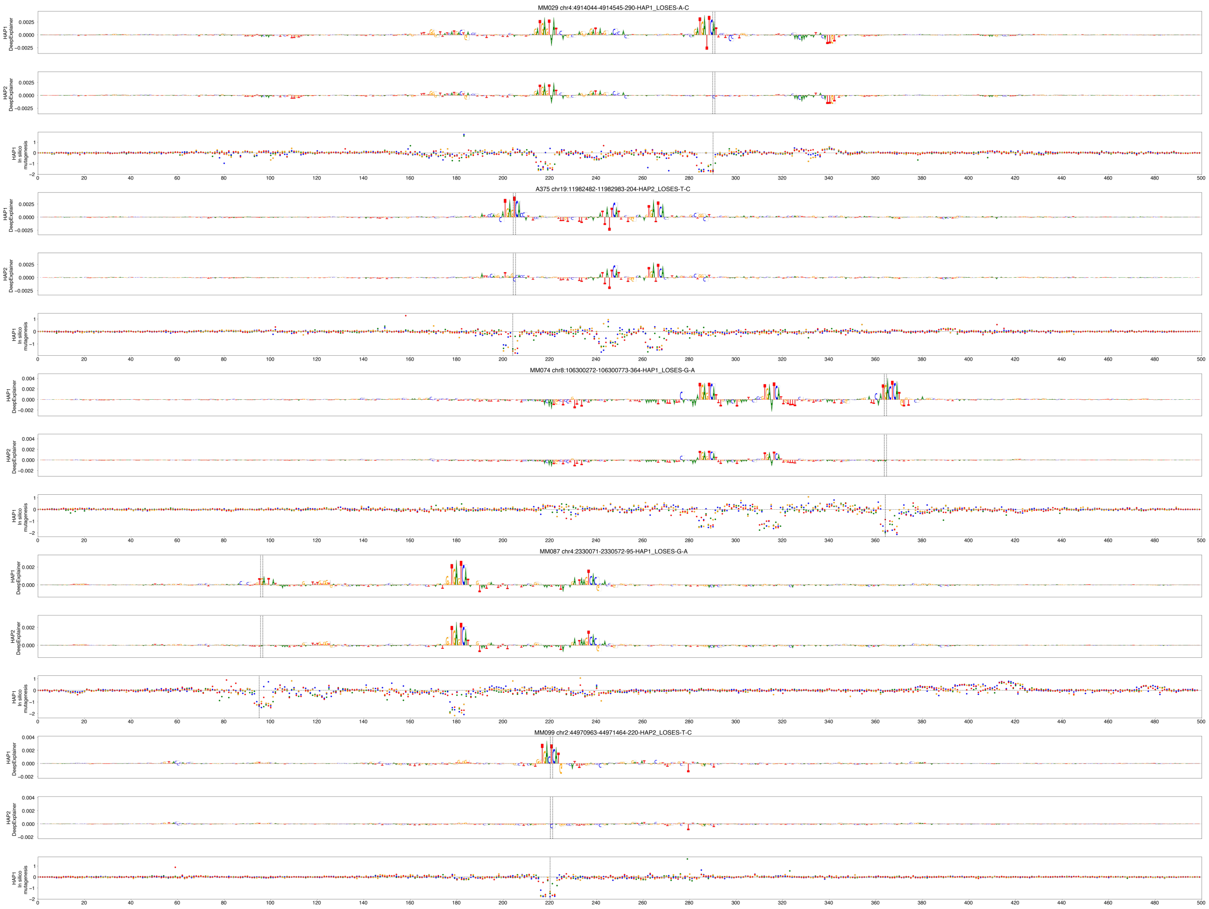


**Supplementary Figure 11.** DeepExplainer plots for several examples of MES enhancers in 5 different MM lines (MM029, A375, MM074, MM087, and MM099) with an ASCAV altering an AP-1 binding site.

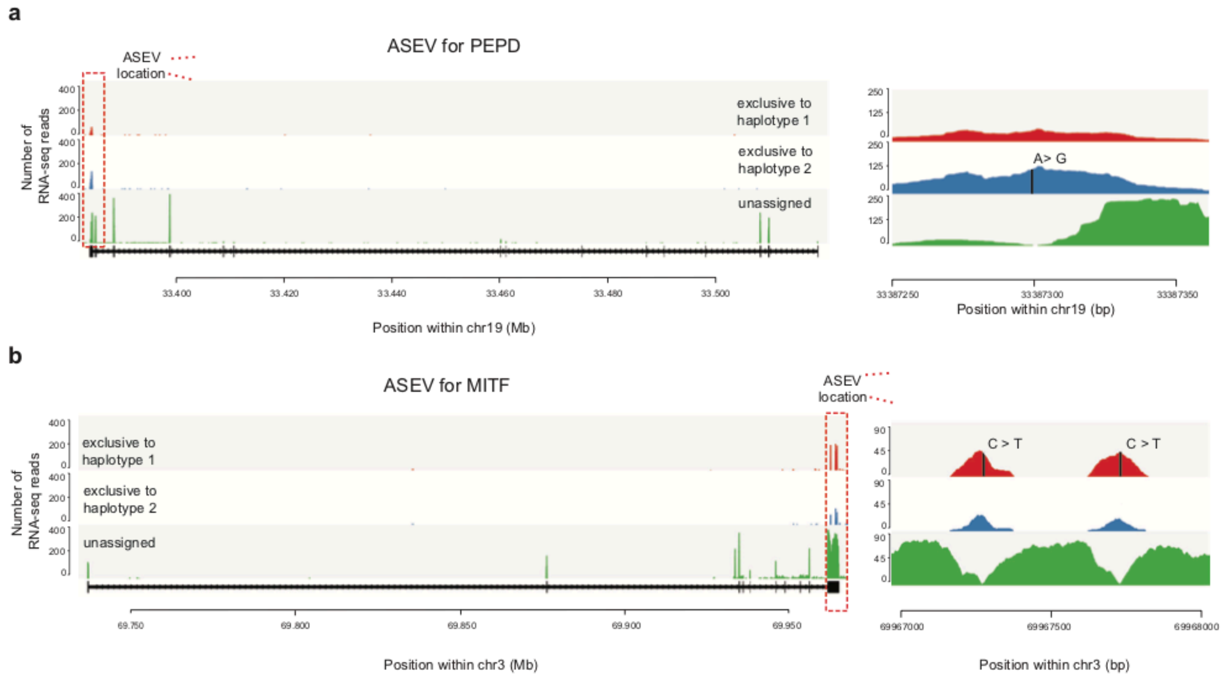


**Supplementary Figure 12.** Haplotype resolved visualization of allele-specific expression (ASE) variants at PEPD and MITF loci.

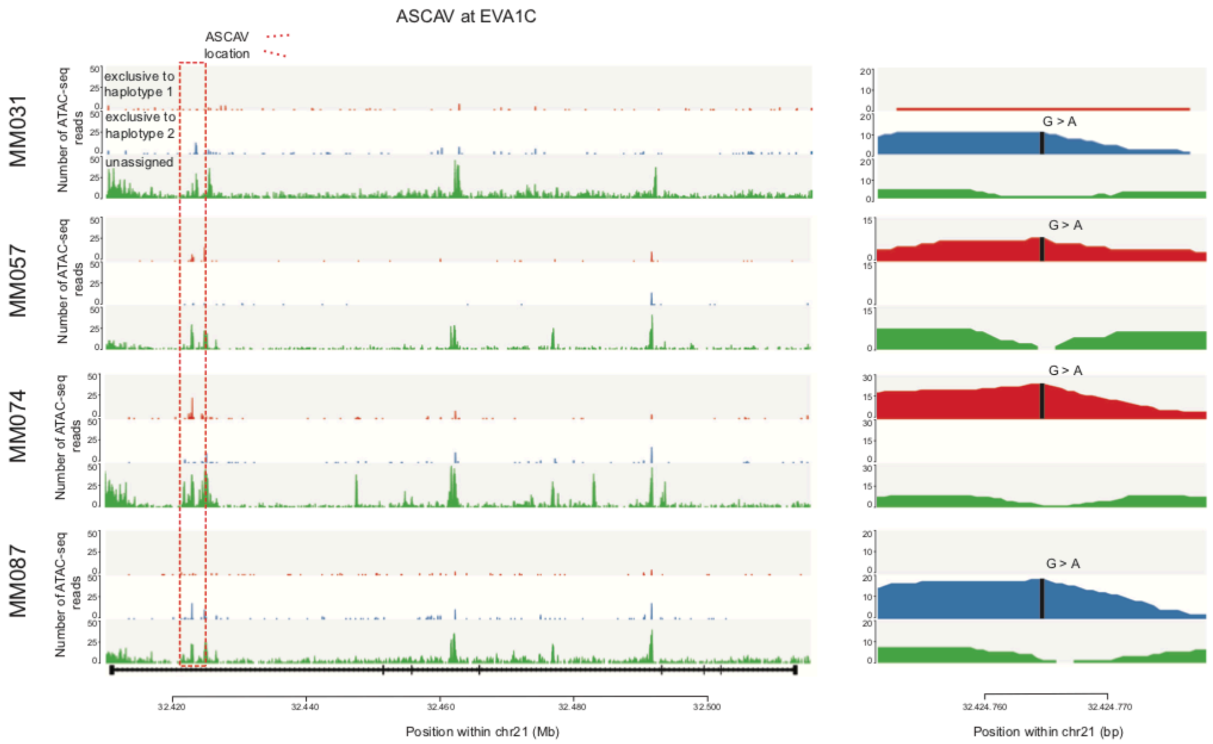


**Supplementary Figure 13.** Haplotype resolved visualization of EVA1C across four melanoma lines with rs2833812. The SNP in the first intron of EVA1C results in allelic imbalance in all lines where the SNP is present.
